# A Novel Hypothalamic Factor, Neurosecretory Protein GM, Causes Fat Deposition in Chicks

**DOI:** 10.1101/2021.07.24.453660

**Authors:** Masaki Kato, Eiko Iwakoshi-Ukena, Megumi Furumitsu, Kazuyoshi Ukena

**Affiliations:** Laboratory of Neurometabolism, Graduate School of Integrated Sciences for Life, Hiroshima University, Higashi-Hiroshima, Hiroshima 739-8521, Japan

**Keywords:** neurosecretory protein, chicken, hypothalamus, fat deposition, chronic intracerebroventricular infusion

## Abstract

We recently discovered a novel cDNA encoding the precursor of a small secretory protein, neurosecretory protein GM (NPGM), in the mediobasal hypothalamus of chickens. Although our previous study showed that subcutaneous infusion of NPGM for 6 days increased body mass in chicks, the chronic effect of intracerebroventricular (i.c.v.) infusion of NPGM remains unknown. In this study, chronic i.c.v. infusion of NPGM for 2 weeks significantly increased body mass, water intake, and the mass of abdominal fat in chicks, whereas NPGM did not affect food intake, muscle mass, or blood glucose concentration. Morphological analyses revealed that fat accumulation occurred in both liver and abdominal fat. These results indicate that NPGM may participate in fat storage in chicks.

## INTRODUCTION

Energy intake through feeding behavior is a necessary element for the maintenance of animal life and growth. However, excessive food intake can cause diseases, such as obesity (Moore et al., 2014). In general, it is known that animals have a complex endocrine system that controls their appetite to avoid obesity (Lawrence et al., 1999; Geary, 2000). Poultry, especially chickens, are an important agricultural species and have been repeatedly selected for meat and/or egg production, with emphasis on feed efficiency and growth rate. However, there is concern that excessive food intake and fat accumulation in broiler chickens can lead to reduced growth rates and metabolic diseases, resulting in lower food production (Knowles et al., 2008). This problem is undesirable from the standpoint of animal health and production efficiency. Therefore, it is important to control growth and fat accumulation in chickens to enable efficient meat production. Understanding the mechanisms of fat accumulation in birds at the molecular and cellular levels will provide sufficient knowledge to improve these problems. In contrast, chicken fat is also one of healthy foods (Peña-Saldarriaga et al., 2020). Several studies have shown that the nutrient composition of diets affects adipose tissue development and lipid metabolism in chickens (Wang et al., 2017). However, the contribution of hormones to lipid metabolism in chickens and other birds has not yet been revealed. Therefore, it is likely that unknown factors are involved in these processes in avian species.

Recently, we discovered a novel cDNA encoding the precursor of a small secretory protein in the chick hypothalamus (Ukena et al., 2014; Shikano et al., 2018a). We named the small protein of the 83-amino acid residues as neurosecretory protein GM (NPGM) (Ukena et al., 2014; Shikano et al., 2018a). The NPGM gene is highly conserved in vertebrates, including chickens, rats, and humans (Ukena et al., 2014). Our previous studies identified the localization of NPGM in the chick brain by in situ hybridization and immunostaining. NPGM-producing cells are localized in the infundibular nucleus (IN) and the medial mammillary nucleus (MM) of the hypothalamus in chicks (Shikano et al., 2018a). These nuclei are known to be feeding and metabolic centers in chicks. Furthermore, we found that the expression levels of *NPGM* mRNA gradually decreased during post-hatching development (Shikano et al., 2018a). These results suggest that NPGM is a novel hypothalamic factor involved in energy metabolism immediately after hatching in chicks. Subsequently, we performed chronic subcutaneous infusion to clarify the physiological functions of NPGM in chicks. Chronic subcutaneous infusion of NPGM for 6 days increased the body mass gain in chicks (Shikano et al., 2018b). It was not clear which parts of the body gained mass, although NPGM seemed to induce the largest increase in abdominal fat mass (Shikano et al., 2018b). In the present study, we analyzed the effects of chronic intracerebroventricular (i.c.v.) infusion of NPGM for 2 weeks on food intake, water intake, and body composition, including fat accumulation and lipid metabolism in abdominal fat and liver.

## MATERIALS AND METHODS

### Animals

One-day-old male layer chicks were purchased from a commercial company (Nihon layer, Gifu, Japan) and housed in a windowless room at 28 °C on a 20–hour light (4:00–24:00) /4–hour dark (0:00–4:00) cycle. The chicks had *ad libitum* access to food and water.

### Production of Chicken NPGM

Chicken NPGM was synthesized using fluorenylmethyloxycarbonyl (Fmoc) and a peptide synthesizer (Syro Wave; Biotage, Uppsala, Sweden) according to our previous method (Masuda et al., 2015; Shikano et al., 2019). The purity of the protein was >95%. Lyophilized NPGM was weighed using an analytical and precision balance (AP125WD; Shimadzu, Kyoto, Japan).

### Chronic I.C.V. Infusion of Chicken NPGM for 2 Weeks

Chronic i.c.v. infusion of chicken NPGM was performed according to a previously reported method (Shikano et al., 2018c; Shikano et al., 2019). Eight-day-old chicks were i.c.v-infused with 0 (vehicle control) or 15 nmol/day NPGM. The dose was determined based on previous studies (Shikano et al., 2018c; Shikano et al., 2019; Rich et al., 2007). Body mass, food intake, and water intake were measured daily (between 9:00 and 10:00) throughout the experiment. After 2 weeks of chronic i.c.v. infusion of NPGM, chicks were sacrificed by decapitation, and the masses of the liver, abdominal fat, subcutaneous fat, pectoralis major muscle, pectoralis minor muscle, and biceps femoris muscle were measured. Blood samples were taken at the endpoint and centrifuged at 800×g for 15 min at 4°C to separate the serum. The blood glucose levels were measured using a GLUCOCARD G+ meter (Arkray, Kyoto, Japan).

### Morphological Analyses

Oil Red O staining of the livers and hematoxylin and eosin staining of the abdominal fat were performed according to a previously reported method (Shikano et al., 2019).

### Real-Time PCR

RNA extraction was performed as previously described (Shikano et al., 2019). PCR amplifications were performed using THUNDERBIRD SYBR qPCR Mix (TOYOBO, Osaka, Japan): 95°C for 20 s followed by 40 cycles at 95°C for 3 s and 60°C for 30 s using a real-time thermal cycler (CFX Connect; BioRad, Hercules, CA).

The amplification of the lipid metabolic factors was performed using the primer sets listed in Supplementary Table 1. Acetyl-CoA carboxylase (*ACC*), fatty acid synthase (*FAS*), stearoyl-CoA desaturase 1 (*SCD1*), malic enzyme (*ME*), peroxisome proliferator-activated receptor γ (*PPARγ*), and fatty acid transporter 1 (*FATP1*) are lipogenic enzymes and related factors, and peroxisome proliferator-activated receptor *α* (*PPARα*), carnitine palmitoyltransferase 1a (*CPT1a*), lipoprotein lipase (*LPL*), adipose triglyceride lipase (*ATGL*), and comparative gene identification-58 (*CGI-58*) are lipolytic enzymes and related factors.

Amplification of feeding, water intake, and growth-related factors was performed using the primer sets listed in Supplementary Table 1. We chose neurosecretory protein GL (*NPGL*) as a paralogous gene of NPGM; neuropeptide Y (*NPY*) and agouti-related peptide (*AGRP*) as orexigenic factors; pro-opiomelanocortin (*POMC*), glucagon-like peptide-1 (*GLP-1*), and cholecystokinin (*CCK*) as anorexigenic factors; and angiotensinogen (*AGT*) and angiotensin-converting enzyme (*ACE*) as water intake-related factors in the hypothalamus. In the pituitary gland, we chose growth hormone (*GH*), prolactin (*PRL*), thyroid-stimulating hormone (*TSH*), and *POMC*. The relative quantification for each expression was determined by the 2^−ΔΔCt^ method using β-actin (*ACTB*) as an internal control.

### Statistical Analysis

Data were analyzed with Student’s *t*-test for endpoint body mass, cumulative food intake, cumulative water intake, tissue mass, blood glucose level, and mRNA expression or two-way repeated-measures analysis of variance (ANOVA) followed by Bonferroni’s test for body mass gain and daily food and water intake. The significance level was set at *P* < 0.05. All results are expressed as the mean ± SEM.

## RESULT

### Effects of Chronic I.C.V. Infusion of NPGM on Body Mass, Food Intake, Water Intake

To investigate the effect of chronic i.c.v. infusion of NPGM on energy metabolism, we measured body mass, food intake, and water intake for 2 weeks. The results showed that chronic infusion of NPGM significantly increased daily body mass gain and final body mass after 2 weeks (Figure 1A and B). However, daily and cumulative food intake remained unchanged (Figure 1C and D). The daily water intake was almost unchanged, but the cumulative water intake increased after 2 weeks of NPGM infusion (Figure 1E and F).

**Figure 1.**
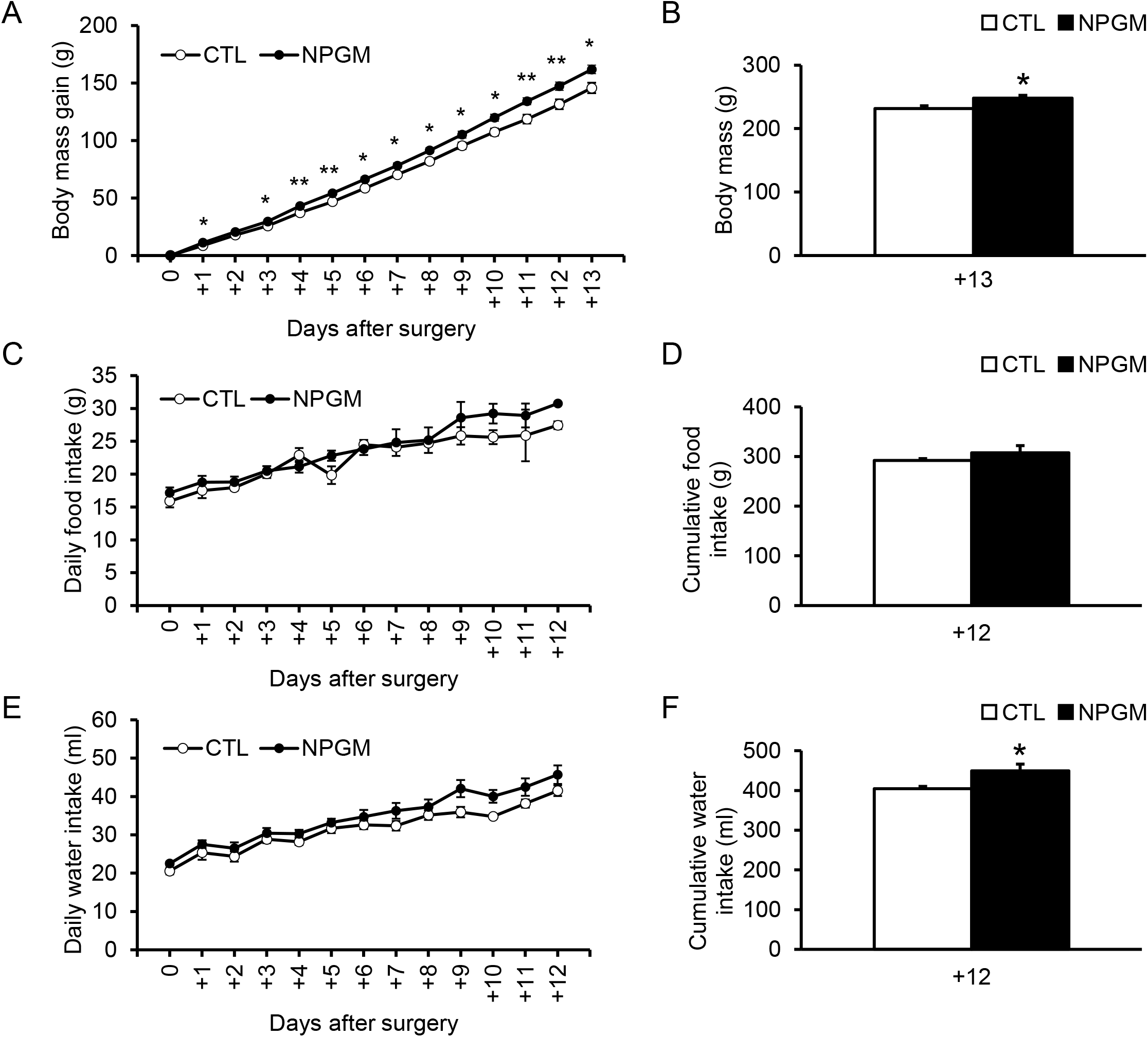
Effect of chronic i.c.v. infusion of NPGM on body mass gain, food intake, and water intake. The results were obtained by the infusion of the vehicle (control; CTL) and NPGM. The change in the body mass gain after surgery (A, B). The daily and cumulative food intake (C, D). The daily and cumulative water intake (E, F). Data are expressed as the mean ± SEM (n = 7–8). Data were analyzed using the student *t*-test and two-way repeated-measures analysis of variance (ANOVA). An asterisk indicates a statistically significant difference (\**P* < 0.05, \*\**P* < 0.01).

### Effects of Chronic I.C.V. Infusion of NPGM on Body Composition

When we analyzed the effect of NPGM on body composition and blood glucose level, chronic infusion of NPGM increased the mass of the abdominal fat and tended to increase the masses of the liver and subcutaneous fat (Figure 2A and B). On the other hand, no change was observed in the muscle masses of the pectoralis major and pectoralis minor muscles after the infusion of NPGM, and a slight increase was found in the mass of the biceps femoris muscle (Figure 2C). Furthermore, the blood glucose levels were not altered by NPGM (Figure 2D).

**Figure 2.**
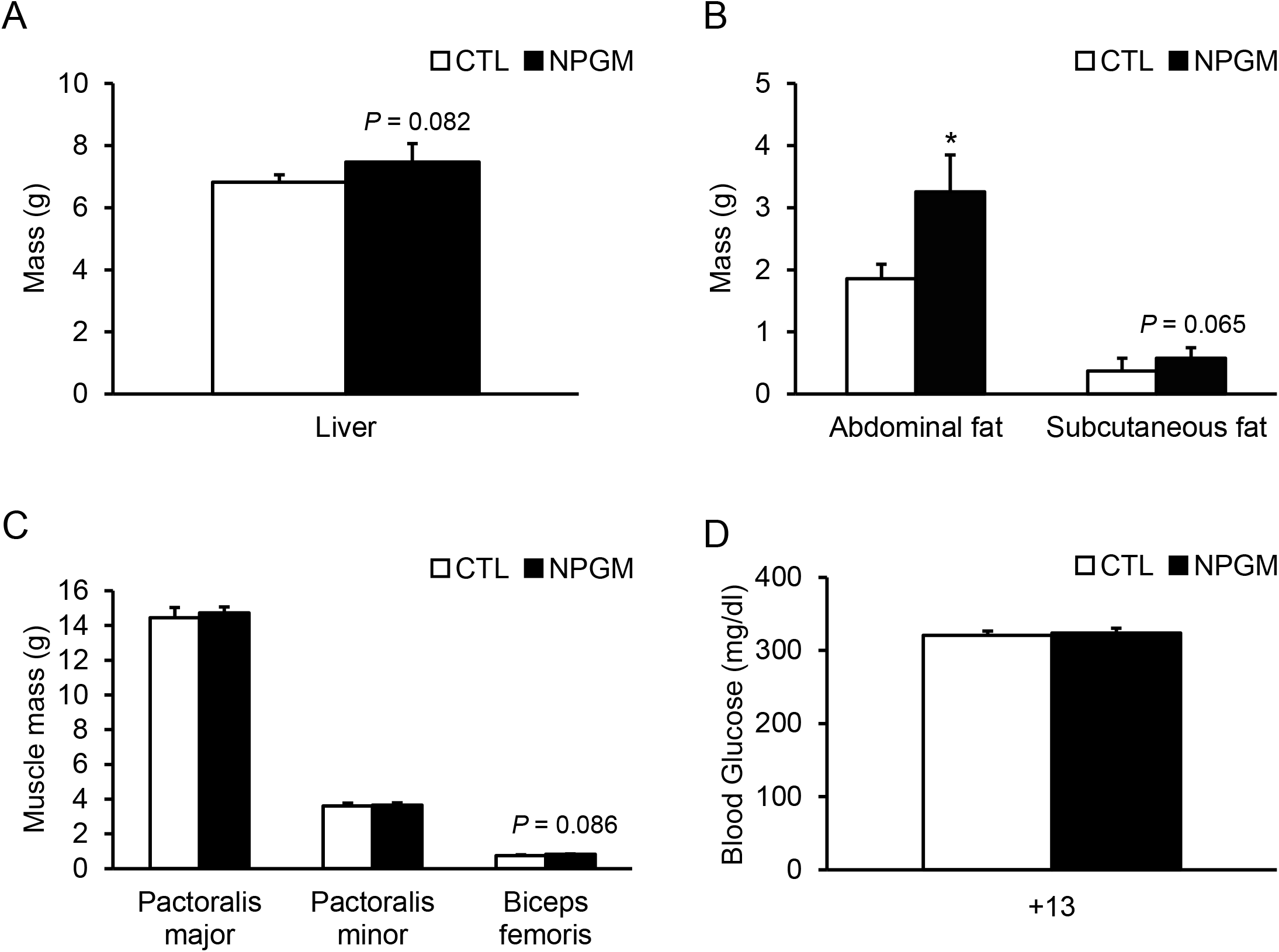
Effect of chronic i.c.v. infusion of NPGM on body composition. The results were obtained by the infusion of the vehicle (control; CTL) and NPGM for 2 weeks. The mass of the liver (A), abdominal fat, and subcutaneous fat (B), pectoralis major muscle, pectoralis minor muscle, and biceps femoris muscle (C), and blood glucose level (D) were measured at the end of the experiment. Data are expressed as the mean ± SEM (n = 7–8). Data were analyzed by the Student’s *t*-test. Asterisks indicate statistically significant differences (\**P* < 0.05).

### Effects of Chronic I.C.V. Infusion of NPGM on Lipid Deposition in the Liver and Abdominal Fat

To elucidate the cause of the increase in the masses of the liver and adipose tissues, we elucidated the effect of fat accumulation by morphological analyses. Oil Red O staining of the liver demonstrated that lipid droplets were deposited after chronic i.c.v. infusion of NPGM (Figure 3A).

**Figure 3.**
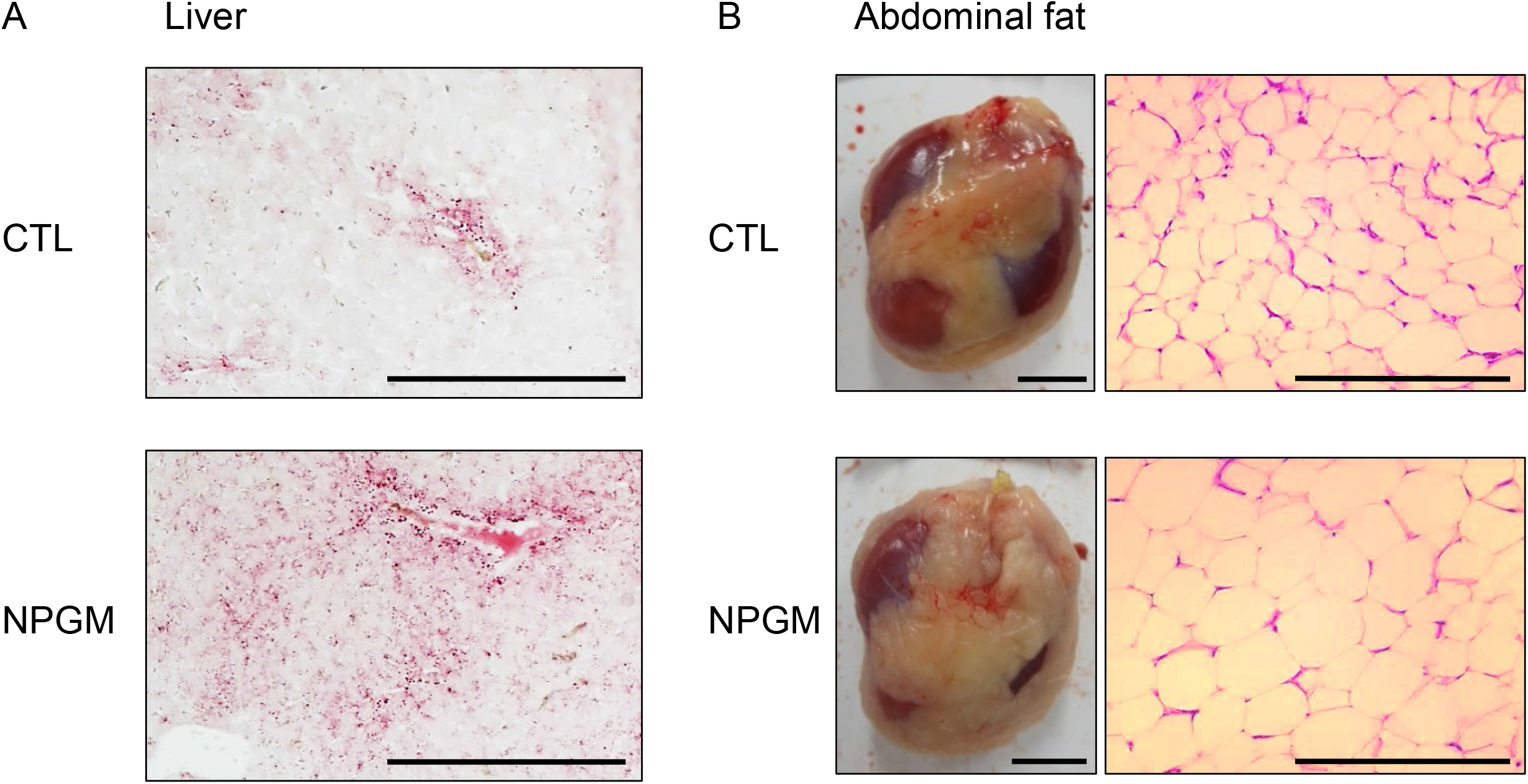
Effect of chronic i.c.v. infusion of NPGM on lipid deposition in the liver and lipid droplets in the abdominal fat. The results were obtained 2 weeks after infusion of the vehicle (control; CTL) and NPGM. In the liver, representative photomicrographs of the sections were stained by Oil Red O (scale bar = 100 μm) (A). For the abdominal fat, exterior photographs of the abdominal fat around the gizzard (left panel, scale bar = 1 cm) and representative photographs of the sections stained by hematoxylin and eosin (right panel, scale bar = 100 μm) (B) were obtained.

We investigated the abdominal fat and found an increase in abdominal fat around the gizzard in NPGM-treated chicks compared with controls (Figure 3B, lower left panel). Hematoxylin-eosin staining revealed that the adipocytes in the abdominal fat of the NPGM-treated chicks were larger than those of the control group (Figure 3B, lower right panel).

### Effects of Chronic I.C.V. Infusion of NPGM on the mRNA Expression of Lipogenic and Lipolytic Factors in the Liver and Abdominal Fat

To explore the molecular mechanism of fat accumulation in the liver and abdominal fat by chronic i.c.v. infusion of NPGM, we investigated the mRNA expression levels of the lipid metabolic factors. We analyzed the mRNA expression levels of *ACC, FAS, SCD1, ME, PPARγ, FATP1, PPARα, CPT1a, LPL, ATGL, and CGI-58* after chronic infusion of NPGM. In the liver, NPGM decreased the mRNA expression of *PPARα* (Figure 4A), whereas in abdominal fat, the mRNA expression levels did not change (Figure 4B).

**Figure 4.**
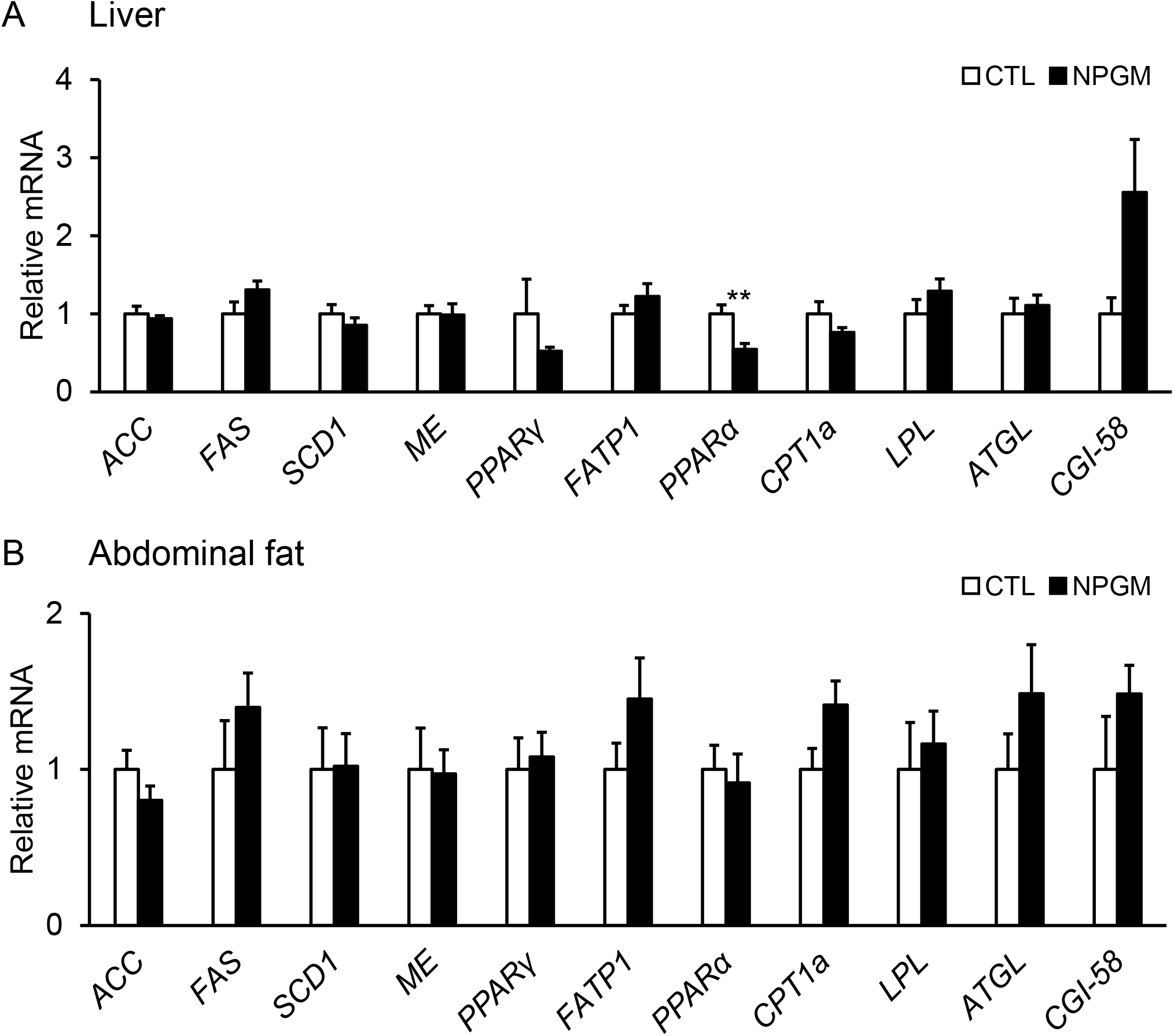
Effect of chronic i.c.v. infusion of NPGM on the mRNA expression of lipogenic and lipolytic factors [acetyl-CoA carboxylase (*ACC*), fatty acid synthase (*FAS)*, stearoyl-CoA desaturase 1 (*SCD1*), malic enzyme (*ME*), peroxisome proliferator-activated receptor γ (*PPARγ*), fatty acid transporter 1 (*FATP1*), peroxisome proliferator-activated receptor α (*PPARα*), carnitine palmitoyltransferase 1a (*CPT1a*), lipoprotein lipase (*LPL*), adipose triglyceride lipase (*ATGL*), and comparative gene identification-58 (*CGI-58*)] in the liver (A) and abdominal fat (B). The results were obtained 2 weeks after infusion of the vehicle (control; CTL) and NPGM. Data are expressed as the mean ± SEM (n = 7–8). Data were analyzed by Student’s *t*-test. An asterisk indicates a statistically significant difference (\*\**P* < 0.01).

### Effects of Chronic I.C.V. Infusion of NPGM on the mRNA Expression for Feeding, Water Intake, and Pituitary Hormones in the Hypothalamus and Pituitary Gland

Based on the results of body mass gain and water intake, we investigated the mRNA expression levels of the endocrine hormones and related factors, which are involved in feeding, water intake, and growth in the hypothalamus and pituitary gland after chronic infusion of NPGM. NPGM did not change the mRNA expression levels in the hypothalamus (Supplementary Figure 1A), whereas NPGM decreased the expression of *PRL* mRNA in the pituitary gland (Supplementary Figure 1B).

## DISCUSSION

In this study, we investigated the effects of a novel small protein, NPGM, on the body composition of chicks by chronic i.c.v. infusion for 2 weeks. Our data showed that the infusion of NPGM increased body mass and cumulative water intake without changes in food intake. Furthermore, morphological analyses showed that NPGM induced fat accumulation in the liver and abdominal fat. This study is the first to show that NPGM induces fat accumulation in animals.

Several studies have been conducted on fat accumulation in mammals (Peirce et al., 2014; Gesta et al., 2007). However, the mechanism of lipid metabolism in birds has not been established. The biological structures of birds differ from those of mammals, and their endocrine actions are unique. For example, birds do not have heat-producing brown adipose tissue (BAT) and have only white adipose tissue (WAT) (Johnston, 1971). Moreover, birds are hyperglycemic animals that exhibit insulin resistance despite normal insulin activity (Dupont et al., 2008). In chickens, unlike mammals, insulin has little effect on glucose uptake in adipose tissue and does not inhibit lipolysis (Tokushima et al., 2005). Furthermore, glucagon is thought to be the main lipolytic hormone (Scanes et al., 1994). In mammals, leptin is a hormone secreted by adipocytes and acts on the hypothalamus to suppress eating and obesity (Halaas et al., 1995). Although leptin and leptin receptors have been studied extensively in mammals and have been applied to humans (Friedman and Halaas, 1998), it has long been debated whether leptin is conserved in birds (Taouis et al., 1998; Friedman-Einat et al., 1999). In addition, the presence of leptin in the peripheral circulation has been questioned because it was not detected in the blood of birds (Seroussi et al., 2016). Recently, the leptin gene has been identified in some birds, and avian leptin is expressed in the digestive organs (Friedman-Einat and Seroussi, 2019; Seroussi et al., 2017; Beauclair et al., 2019; Seroussi et al., 2019). These findings suggest that avian leptin may have a different physiological function from that of lipid metabolism in mammals. The present study showed that NPGM is a new player in regulating fat deposition in avian species, chicken.

We have also reported a paralog gene with NPGM, which is named neurosecretory protein GL (NPGL) in vertebrates (Ukena et al., 2014; Ukena, 2018). The NPGL gene is also highly conserved in vertebrates, including chickens, rats, mice, and humans (Ukena et al., 2014; Ukena, 2018). Sequence similarity between mature NPGM and NPGL is 54% in chickens (Shikano et al., 2018b). Our previous studies showed that NPGL-immunoreactive cells were localized in the IN and MM of the hypothalamus, and parts of NPGL-immunoreactive cells were identical to NPGM-immunoreactive cells in the MM in 1- and 15-day-old chicks and the IN in 15-day-old chicks (Shikano et al., 2018a). The expression levels of *NPGL* mRNA gradually increased during post-hatching development. In contrast, the expression level of *NPGM* mRNA gradually decreased after hatching, suggesting that NPGM and NPGL have different timings of action during the growth process (Shikano et al., 2018a). Functional analysis of NPGL has already been demonstrated in rats, mice, and chicks (Iwakoshi-Ukena et al., 2017; Fukumura et al., 2021; Matsuura et al., 2017; Shikano et al., 2020; Shikano et al., 2018c; Shikano et al., 2019). The results showed that the long-term administration of NPGL stimulated food intake and induced fat accumulation in rats, mice, and chicks. In contrast, NPGM treatment did not change food intake or the gene expression levels of feeding and growth regulators (*NPY, AgRP, POMC, GLP-1, CCK, GH*, and *TSH*) in the hypothalamus and pituitary gland in this study. Therefore, it is likely that NPGM does not directly regulate feeding behavior in chicks. In addition, the expression levels of water intake-regulating factors (*AGT* and *ACE*) did not change, although the amount of drinking water was increased by NPGM infusion. Therefore, NPGM does not seem to be a strong factor that regulates drinking behavior.

In our previous studies, NPGL increased the mass of WAT through *de novo* lipogenesis in rats and chicks, resulting in fast body mass gain (Iwakoshi-Ukena et al., 2017; Shikano et al., 2019). In general, birds are known to use the liver rather than adipose tissue as the primary site of *de novo* lipogenesis (Leveille et al., 1975). Glucose taken up by the liver is converted into fatty acids and cholesterol and, subsequently, fatty acids are converted into triacylglycerols (TAGs). Cholesterol and TAGs are transported by very low-density lipoproteins (VLDL) to the adipose tissue where they are stored (Hermier, 1997). However, we found that *de novo* lipogenesis by NPGL in chicks occurred in the adipose tissue but not in the liver (Shikano et al., 2019). This result indicates that NPGL is a neuropeptide that upregulates *de novo* lipogenesis in the adipose tissues of chicks. In the present study, chronic i.c.v. infusion of NPGM induced body mass gain in chicks, similar to NPGL. However, mRNA expression analysis of lipid metabolism factors involved in lipogenesis and lipolysis showed little change in the expression levels of lipid metabolism factors in the liver and adipose tissue, except for a decrease in the lipolytic factor PPARα in the liver. These results suggest that *de novo* lipogenesis by NPGM does not occur in these tissues. Therefore, the mechanism of action of NPGM and NPGL for fat accumulation may differ in the same animals. It is also known that lipogenesis and lipolysis in the adipose tissue are controlled by the hypothalamus via the sympathetic nervous system in mammals (Buettner et al., 2008; Scherer et al., 2011). Therefore, NPGM may be involved in the sympathetic control of fat accumulation in chickens. Future studies are needed to classify the target regions and neural networks of NPGM-producing neurons in chickens.

In this study, *PRL* mRNA expression was decreased in the pituitary gland after NPGM infusion. PRL is known to be involved in more than 300 actions in vertebrates (Bole-Feysot et al., 1998). In birds, the most studied role of PRL is involved in actions during the egg incubation phase (March et al., 1994). In addition, PRL has been reported to increase feeding behavior and activate LPL in adipocytes (Das, 1991; Garrison and Scow, 1975). In the present study, there was no change in food intake or *LPL* mRNA expression levels after NPGM infusion. However, NPGM may inhibit the increase in food intake and the activation of lipolysis through the suppression of PRL expression. In the future, it will be necessary to clarify the exact relationship between NPGM and PRL in chickens.

The present data on NPGM-induced fat accumulation in chickens may facilitate future applications in animal agriculture, such as controlling fat accumulation in poultry. The French food, *foie gras*, is a fatty liver, and the artificial production of *foie gras* is a problem related to animal welfare (Litt et al., 2020). In addition, chicken fat is becoming a healthy food because of the high amounts of unsaturated fatty acids (Peña-Saldarriaga et al., 2020). The application of NPGM to fat deposition in the liver and abdominal fat will have an impact on agricultural and livestock industries.

To clarify the exact role of NPGM action in the brain, the receptor for NPGM needs to be discovered. However, to date, receptors for NPGM and NPGL have not yet been discovered in vertebrates. It is known that fat accumulation is essential for birds, not only to sustain life but also to maintain bird-specific behaviors such as migration (Guglielmo. 2018; Price. 2010). Therefore, NPGM is considered an important factor in avian survival and reproduction. In the future, comparative analysis using other avian species and other vertebrates will help clarify the physiological role of NPGM in animals.

## DATA AVAILABILITY STATEMENT

The original contributions presented in the study are included in the article/supplementary material, further inquiries can be directed to the corresponding author.

## ETHICS STATEMENT

The experimental protocols were in accordance with the Guide for the Care and Use of Laboratory Animals prepared by Hiroshima University (Higashi-Hiroshima, Japan).

## AUTHOR CONTRIBUTIONS

Conceptualization, M.K. and K.U.; methodology, M.K., E.I-U., M.F. and K.U.; investigation, M.K., E.I-U., M.F. and K.U.; writing—original draft preparation, M.K.; writing—review and editing, M.K. and K.U.; visualization, M.K.; project administration, K.U.; funding acquisition, E.I.-U. and K.U. All authors have read and agreed to the published version of the manuscript.

## ACKNOWKEDGMENTS

We are grateful to Dr. Kenshiro Shikano, Mr. Yuki Narimatsu, and Dr. Keisuke Fukumura for the experimental support.

---

**Supplementary Figure 1**

Effect of chronic i.c.v. infusion of NPGM on the mRNA expression of neurosecretory protein GL (*NPGL*), neuropeptide Y (*NPY*), agouti-related peptide (*AGRP*), pro-opiomelanocortin (*POMC*), glucagon-like peptide-1 (*GLP-1*), cholecystokinin (*CCK*), angiotensinogen (*AGT*), and angiotensin-converting enzyme (*ACE*) in the hypothalamus (A). In the pituitary gland, we chose growth hormone (*GH*), prolactin (*PRL*), thyroid-stimulating hormone (*TSH*), and *POMC* (B). The results were obtained 2 weeks after infusion of the vehicle (control; CTL) and NPGM. Data are expressed as the mean ± SEM (n = 7–8). Data were analyzed by Student’s *t*-test. An asterisk indicates a statistically significant difference (\*\*\**P* < 0.005).

