## Supplementary Figure and Table for "A Novel Hypothalamic Factor, Neurosecretory Protein GM, Causes Fat Deposition in Chicks"

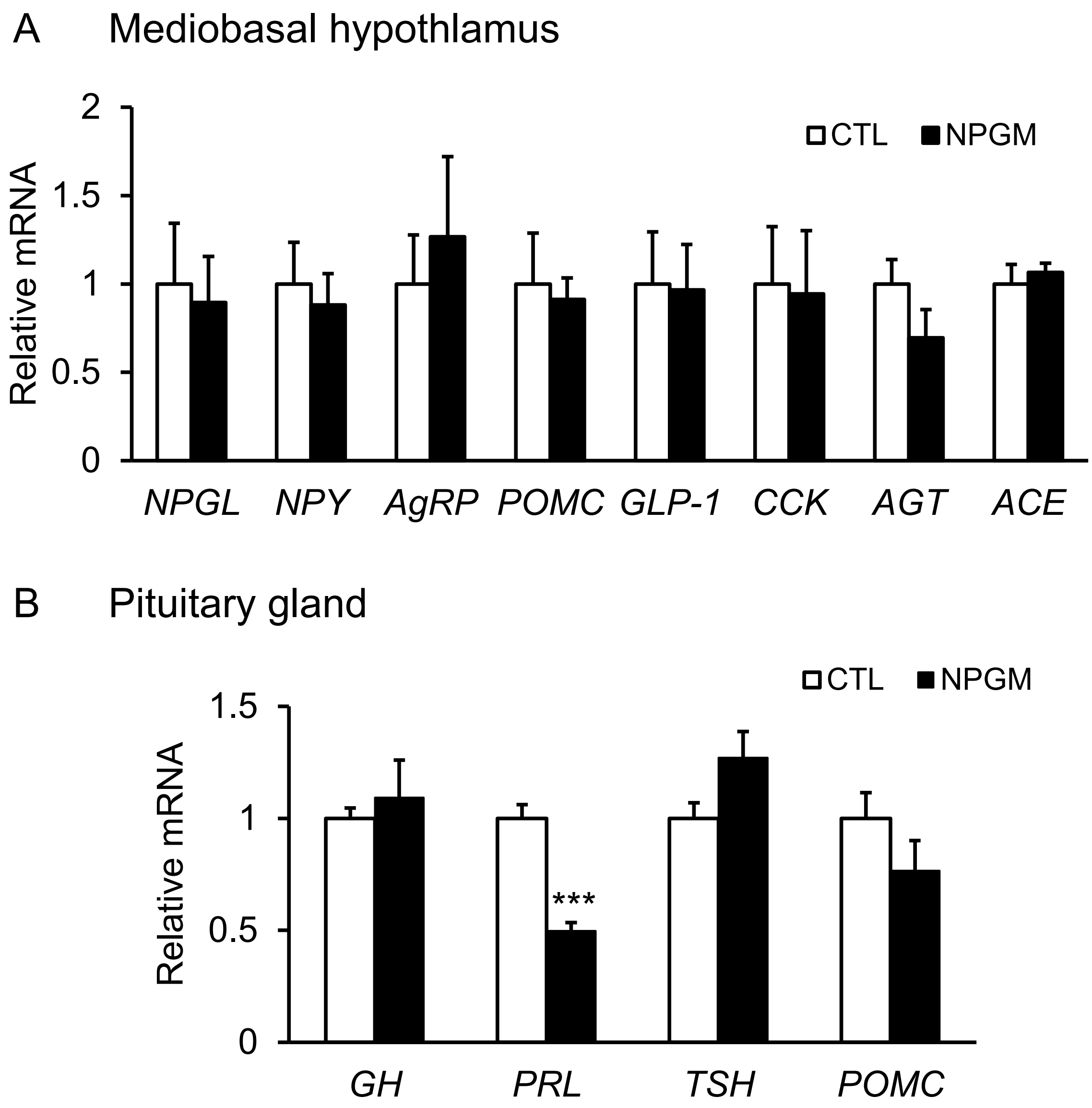


Supplementary Figure 1

Table 1. Sequences of oligonucleotide primers for real-time PCR

| Gene | Forward primer | Reverse primer | Accession no. |
|---|---|---|---|
| *ACC* | AATGGCAGCTTTGGAGGTGT | TCTGTTTGGGTGGGAGGTG | NM_205505.1 |
| *FAS* | CCAACGATTACCCGTCTCAA | CAGGCTCTGTATGCTGTCCAA | NM_205155.2 |
| *SCD1* | AGTGGTGTTGCTGTGCTTCA | CTAAGGTGTAGCGCAGGATG | NM_204890.1 |
| *ME* | AGTGCCTACCTGTGATGTTG | GGCTTGACCTCTGATTCTCT | NM_204303.1 |
| *PPARγ* | TCAAGCATTTCTTCACCACACT | ATTGCACTTTGGCAATCCTGG | NM_001001460.1 |
| *FATP1* | TACAATGTGCTCCAGAAGGG | GTCTGGTTGAGGATGTGACTC | NM_001039602.2 |
| *PPARα* | TGCTGTGGAGATCGTCCTGGT | AGAGGAAGATATCGTCAGGATGG | NM_001001464.1 |
| *CPT1a* | TGATCTGAAGAAGAACCCTGAGAT | TCCAAAGCGATGAGAATCCG | NM_001012898.1 |
| *LPL* | CAGTGCAACTTCAACCATACCA | AACCAGCCAGTCCACAACAA | NM_205282.1 |
| *ATGL* | CACTGCCATGATGGTCCCCTA | CCACAAGGAGATGCTGAAGAA | NM_001113291.1 |
| *CGI-58* | ACCGTGGTTTATGGAGCACG | GAAACAGTGTGCAAACAGAGCC | NM_001278145.1 |
| *NPGL* | CTAGGAAAAAGACAGCTTGC | CTTTCTTCGTCAGAACTGGT | NM_001389496 |
| *NPY* | ACATGGCCAGATACTACTCG | ACAAGAGGTCTGAGATCAGTG | NM_205473.1 |
| *AgRP* | AGGCCAGACTTGGATCAGATG | ACTCCAGGAGGCGGACAC | NM_001031457.1 |
| *POMC* | AGGAGTCGGCTGAGAGTTA | TTCCTCTTCCTCCTCTTCTT | NM_001031098.1 |
| *GLP-1* | CGTCATTCACAAGGCACATTC | GTCATTCTCTTTGTCCTCCTGTCC | NC_052534.1 |
| *CCK* | AGGTTCCACTGGGAGGTTCT | CGCCTGCTGTTCTTTAGGAG | NC_052533.1 |
| *AGT* | AGCAGGTTTGAGAGGCAATGA | GATTCCACCACTTCCCCAGG | NC_052534.1 |
| *ACE* | GCCAAACTCAGGGAGGTGTT | CCCCAGCGTCCATCTTATCC | NC_052558.1 |
| *GH* | CCAGAGTCCATCACAATACC | AGCCAACAGAGAGAAGATGA | NC_052558.1 |
| *PRL* | AAGAAGCTCCAGATACCATTCTCT | GAGAGTAAATTTCATTTCCAGCAT | NC_052533.1 |
| *TSH* | CCACCATCTGCGCTGGAT | GCCCGGAATCAGTGCTGTT | NC_052557.1 |
| *ACTB* | CCAGAGTCCATCACAATACC | AGCCAACAGAGAGAAGATGA | NM_205518.1 |

Supplementary Table 1
